# Tumor necrosis factor receptor 2 inhibits HIV-1 infection by blocking the binding of gp120 to CD4

**DOI:** 10.1101/2025.08.05.668613

**Authors:** Yang Gao, Zhonghao Chen, Yibo Chen, Yang Yang, Yiru Wang, Ci Zhu, Ping Liao, He Song, Xin Chen

**Affiliations:** Institute of Chinese Medical Sciences, State Key Laboratory of Quality Research in Chinese Medicine, University of Macau, Macau SAR, China; Department of Pharmaceutical Sciences, Faculty of Health Sciences, University of Macau, Macau SAR, China; MoE Frontiers Science Center for Precision Oncology, University of Macau, Macau SAR, China

**Author notes:** Address correspondence to: Xin Chen;, He Song. These authors contributed equally to this work (Yang Gao, Zhonghao Chen).

**Keywords:** HIV-1, gp120, TNFR2, TNF, TNFR1, pseudovirus

## Abstract

The HIV-1 envelope glycoprotein gp120 binds to CD4 molecule and then undergoes conformational changes to interact with the co-receptors CCR5 or CXCR4, resulting in cellular entrance. However, certain types of cells, such as macrophages and CD4+Foxp3+ regulatory T cells (Tregs), have shown to resist HIV-1 infection despite co-expressing CD4 and co-receptors. In this study, we found that TNF receptor type II (TNFR2) directly bound to gp120, with the binding site on gp120 in close proximity to that of CD4. Intriguingly, exogenous TNFR2 had the capacity to inhibit the binding of gp120 to CD4 T cells. Furthermore, the infection of CD4^+^CCR5^+^ cells by pseudoviruses containing the HIV-1 envelope was inhibited by TNFR2 protein. In contrast, TNFR1, similar in structure to TNFR2 and sharing the same ligand, failed to inhibit the infection of CD4^+^ T cells by HIV-1 pseudoviruses. This property of TNFR2 may be harnessed in the prevention or treatment of HIV-1 infection and thus warrant future investigation.

**SUMMARY:** This study reveals that TNFR2 directly binds to HIV-1 gp120, inhibiting its interaction with CD4 and reducing HIV-1 infection in CD4+ T cells. The findings highlight TNFR2’s potential role in protecting cells and offer insights for HIV-1 prevention or therapy.

## INTRODUCTION

Human Immunodeficiency Virus-1 (HIV-1), the virus that causes AIDS, infects cells by first binding to the CD4 and then to the co-receptors CCR5 or CXCR4. A gradual decrease in CD4 cells is the characteristic of AIDS, yet many questions relating to the mechanisms underlying this decrease remain unanswered(1). Among all CD4-positive cells, several with high co-receptor expression, such as macrophages and regulatory T cells (Tregs), have been found exhibite reduced susceptibility resistant to HIV-1 infection (2–4). Previous studies have shown that certain subsets of CD4-positive macrophages exhibit relative resistance to HIV-1-induced cytopathic effects and can persist for extended periods during infection(5). Further, it has been repeatedly reported that, during HIV-1 infection, tumor necrosis factor (TNF) down-regulates the expression of CD4 on the surface of macrophages may be the underlying mechanism(2, 6). However, despite the presence of TNF, effector CD4 T cells do not exhibit resistance to HIV, whereas CD4^+^FoxP3^+^ Tregs, have been found to be resistant to HIV-1 infection(4, 7, 8). Interestingly, Tregs could be isolated and expanded *in vitro* from patients infected with HIV-1, demonstrating that functional Tregs derived from the blood and lymphoid tissues of HIV-1 infected individuals maintain their suppressive capacity, thereby indicating that these cells are not intrinsically impaired in the context of HIV infection(9). These suggest that the infection resistance of Tregs and macrophages may be independent of CD4 expression, and the mechanism remains to elucidated.

Previous reports have indicated that TNF and TNF receptors can influence HIV-1 infection(10, 11). For example, it was reported that pre-treatment with TNF mutation selective for TNFR2 delayed the detection of HIV-1 DNA long terminal repeat sequences in macrophages. It has been also shown that, Foxp3, the essential transcription factor for the development and function of Treg cells, markedly inhibited HIV-1 replication when expressed in primary human CD4 T cells(12),(13). Previously, there is compelling evidence that Treg cells predominantly express TNFR2 and the expression of TNFR2 is critical for the sustaining Foxp3 expression in Tregs (14),(15), suggesting that TNFR2 expression and its signaling may be attributable to the resistance of HIV-1 infection by Tregs. Indeed, a virtual molecular docking study suggested that the HIV-1 gp120 could bind to TNFR2(16). In this study, we experimentally verified that TNFR2 protein bond to the HIV-1 envelope protein gp120 with high affinity. Intriguingly, TNFR2 interrupted the interaction between gp120 and CD4. Moreover, we found that TNFR2 could inhibit the infection of CD4^+^CCR5^+^ cells by pseudoviruses containing the HIV-1 envelope. This property of TNFR2 may be therapeutically harnessed, and thus merit further investigation.

## METHODS AND MATERIALS

### Cell culture

The Jurkat cell line and HEK293T cell line were purchased from American Type Culture Collection (Manassas, USA). Jurkat cells were cultured in RPMI-1640 (Gibco, USA) supplemented with 10% FBS. HEK293T cells were cultured in DMEM (Gibco, USA) supplemented with 10% FBS. Freshly isolated PBMCs were obtained from healthy donors through the Macao Blood Transfusion Service after they provided informed consent approved by the institutional review board. Then PBMCs were cultured in the T551 medium (Takara, Japan) containing 10% FBS. The incubation was conducted at 37 ℃ in humidified air containing 5% CO_2_ in a tissue culture flask. The stable cells lines, Jurkat-CCR5, Jurkat-TNFR2-CCR5, 293T-TNFR2, 293T-CD4-CCR5, 293T-TNFR1-CCR5 and 293T-TNFR2-CCR5 were established by lentiviral transduction.

### Reagents

Antibodies were purchased from BD Pharmingen (USA) consisted of PerCP-Cy5.5 anti-human CD4 (Catalog No:566316), APC Streptavidin (Catalog No:554067), PE-Cy7 anti-His (Catalog No: 362620), BV421 anti-human TNFR2 (Catalog No: 562783) were purchased from Biolegend (USA). The human immunodeficiency virus 1 (HIV-1) (group M, subtype B, Isolate MN) gp120 Protein (DNA sequence: AAC31819.1 (Thr30-Arg513), Catalog No: 40405-V08H), human TNFR2 / CD120b protein (DNA sequence: NP_001057.1 (Met1-Asp257), Catalog No: 10417-H08H) were purchased from the Sino Biological (Beijing, China). The human TNFR1 protein (DNA sequence: NP_001056 (Ile 30 - Thr 211) Catalog No: TN1-H5222) was purchased from the ACRO Biosystems (Beijing, China). In addition, TNFR2 protein (DNA sequence: NP_001057 Met1-Gly258) and TNFR1 protein (DNA sequence: NP_001056 Met 1-Thr 211) were also purified and prepared in-house by our laboratory using standard affinity chromatography techniques. CD4 protein (DNA sequence: NP_000607, Catalog No: TP306453) was purchased from OriGene Technologies (Rockville, MD).

### Bio-layer interferometry (BLI)

Biotin labeling of human TNFR2 protein was performed according to the manufacturer’s instructions using a commercial biotinylation kit, followed by purification with a desalting column (Thermo, USA). The concentration of biotinylated TNFR2 was determined using a BCA protein assay kit (Thermo, USA). Biotinylated TNFR2 was diluted in 1× PBS to a final concentration of 5 µg/mL and immobilized onto streptavidin (SA) biosensors (SARTORIUS, Germany, Catalog No: 18-5017) for 600 seconds. Binding kinetics were assessed by exposing the immobilized TNFR2 to recombinant gp120 protein (diluted in PBST: PBS with 0.02% Tween-20, pH 7.4) at concentrations of 2 μM, 1 μM, 0.5 μM, 0.25 μM, and 0.125 μM for 200 seconds (association phase), followed by dissociation in PBST for 300 seconds. PBST alone was used as a reference control to subtract background signal.Data were acquired using an Octet® R2 instrument (SARTORIUS, Germany) and analyzed with ForteBio Data Analysis software version 12.0. A global fitting model based on a 1:1 binding interaction was applied to calculate the association rate constant (kon), dissociation rate constant (koff), and affinity constant (KD = koff/kon).

### Co-immunoprecipitation (Co-IP)

HEK293T cells stably expressing TNFR2 were lysed in ice-cold cell lysis buffer (20 mM Tris-HCl, 150 mM NaCl, 1% Triton X-100, 1 mM EDTA, protease inhibitor cocktail, pH 7.4) and incubated on ice for 30 minutes. The lysate was centrifuged at 14,000 rpm for 15 minutes at 4 °C to remove debris. Supernatants were collected and incubated with recombinant gp120 protein (5 µg/mL final concentration) for 2 hours at 4 °C on a rotator. This mixture was then incubated overnight at 4 °C with 30 µL of Dynabeads Protein G (Thermo Fisher Scientific) pre-conjugated with 2 µg of anti-gp120 antibody (Abcam, USA, Catalog No: ab106578) or control rabbit IgG (Cell Signaling Technology, CST, MA, USA, Catalog No: 2729S). Immunocomplexes were collected by magnetic separation and washed three times with ice-cold PBS containing 0.05% Tween-20. The beads were resuspended in 2× SDS loading buffer and boiled at 95 °C for 5 minutes. Proteins were resolved on 10% Bis-Tris SDS-PAGE gels (Invitrogen) and transferred onto PVDF membranes. Membranes were blocked in 3% BSA in TBST (TBS with 0.1% Tween-20) for 1 hour at room temperature, then incubated overnight at 4 °C with primary anti-TNFR2 antibody (1:1000, CST, USA, Catalog No: 3727). After washing, membranes were incubated with HRP-conjugated secondary antibody (1:5000, CST, Catalog No: 7074) for 1 hour at room temperature and developed using ECL chemiluminescent substrate.

### Enzyme-linked immunosorbent assay (ELISA)

For gp120-TNFR1/TNFR2 binding assays, 96-well high-binding plates were coated with 100 µL of human TNFR1/TNFR2 (5 µg/mL in 1× CBS (15mM Na_2_CO_3_, 35mM NaHCO3, pH 9.6) overnight at 4 °C. After washing with buffer (20 mM Tris-HCl, 500 mM NaCl, 0.05% Tween-20, pH 7.5), wells were blocked with 3% BSA in PBS for 1 hour at room temperature. Serial dilutions of gp120 (1 µM, 0.1 µM, 0.01 µM in blocking buffer) were added and incubated for 2 hours at room temperature. After washing, bound gp120 was detected using anti-His-HRP antibody (1:2000, Thermo, Catalog No: MA1-21315-HRP) and developed with TMB substrate. Reactions were stopped with 1 M H₂SO₄ and absorbance was measured at 450 nm. For competition ELISA, 96-well plates were coated with recombinant CD4 protein (5 µg/mL in CBS) overnight at 4 °C. Mixtures of gp120 (1 µM) and TNFR2 (2 µM) were pre-incubated at room temperature for 30 minutes before being added to the CD4-coated wells and incubated for 2 hours. Binding of gp120 was detected as above using anti-His-HRP.

### Flow Cytometry Assay to Evaluate gp120 Blocking by TNFR2

Jurkat cells and PBMCs were washed and resuspended in FACS buffer (PBS with 2% FBS and 0.1% sodium azide). Cells (1×10^5^ per sample) were incubated with serial dilutions of recombinant TNFR2 protein (800 nM, 400 nM, 200 nM, 100 nM, 50 nM) or TNFR1 (1 µM, 0.6 µM, 0.4 µM) for 1 hour. His-tagged gp120 (5 µg/mL) was then added and incubated for another 1 hour. Cells were washed and stained with PE-Cy7 conjugated anti-His antibody (1:100) for 30 minutes at 4 °C in the dark. For PBMCs, anti-human CD4-PerCP-Cy5.5 antibody (1:100) was used to identify CD4⁺ T cells. After final washes, samples were analyzed using a BD LSRFortessa flow cytometer. Data analysis was performed using FlowJo software.

### HIV-1 pseudovirus infection

The HIV-1 pseudovirus from a CCR5-tropic (R5) strain containing HIV-Env protein encapsulated with green fluorescent protein (GFP)-encoding RNA was obtained from Fubio Biotechnology (Suzhou, Jiangsu, China; Catalog No: FNV5011G). Different concentrations of TNFR2 protein or TNFR1 protein were prepared and incubated with the virus for one hour at 37 °C. Following the incubation period, appropriate target cells were prepared for infection. For infection of 293T-CD4-CCR5 cells, cells were seeded in 24-well plates and maintained under optimal conditions. The pre-treated pseudoviruses were added at a multiplicity of infection (MOI) of 5 or 10. After 16 hours, GFP expression was analyzed by flow cytometry to evaluate infection efficiency. For infection of 293T-CCR5, 293T-TNFR2-CCR5,293T-TNFR1-CCR5 cells, cells were seeded in 24-well plates and maintained under optimal conditions. The pseudoviruses were added at a multiplicity of infection (MOI) of 10. After 16 hours, GFP expression was analyzed by flow cytometry to evaluate infection efficiency.

For Jurkat-CCR5 cell and Jurkat-TNFR2-CCR5 infection, cells were directly infected without prior activation. The virus-protein mixture was added at an MOI of 50, also in the presence of polybrene (1 µg/mL). After 48 hours of incubation, the percentage of GFP-positive cells was measured by flow cytometry to determine infection efficiency.

### The complex structure prediction based on AlphaFold3

Utilized AlphaFold3 to predict the structure of the protein complex(17). Collected the amino acid sequences of TNFR2 (PDB ID: 3ALQ) and gp120 (PDB ID: 4RZ8) from the UniProt database and performed multiple sequence alignments using HHblits. These alignments were then input into AlphaFold3, which was configured for multichain complex predictions. The model generated several candidate structures, which were assessed using the pLDDT scoring system to determine their reliability. The highest-scoring structure was selected as the final model. Visualization and further analysis were conducted using PyMOL(18–20).

### Statistical analysis

All data were presented as means ± SEM, and the statistical analysis was performed by t test or one-way ANOVA test by using GraphPad Prism 10.0. The P value < 0.05 was considered to be statistically significant.

## RESULTS

### HIV-1 envelope protein gp120 interacts with TNFR2

We firstly examined the interaction between the extracellular domain of TNFR2 (residues 1-257) and the gp120 core (residues 30-513) using an Enzyme-Linked Immunosorbent Assay (ELISA). The results showed that at a concentration of 100 nM, the gp120 had the capacity to bind to immobilized TNFR2 (Fig.1A), and this binding occurred in a concentration-dependent manner (Fig.1B), demonstrating a direct interaction between the two proteins. To rule out non-specific binding, we also performed ELISA using uncoated (blank) wells and observed no detectable gp120 signal under the same conditions (Supplementary Fig.1A).

**Figure 1.**
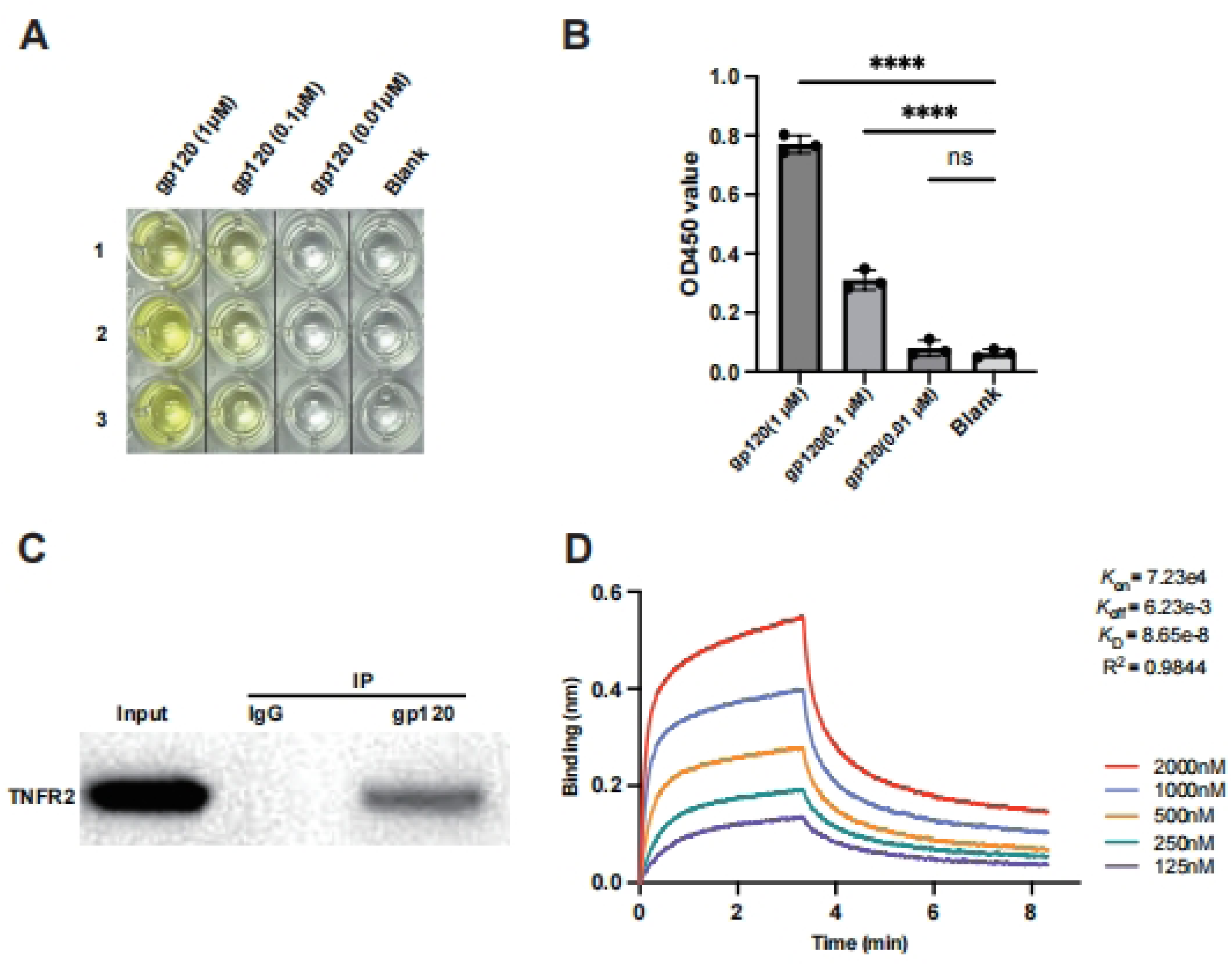
The HIV-1 gp120 protein directly interacts with TNFR2 protein. **(A)** The binding interaction between gp120 and TNFR2 was assessed using an ELISA assay with varying concentrations of gp120 (1μM、0.1μM、0.01μM、blank). The results of color development from the TMB substrate. **(B)** The statistical analysis of the final OD450 measurements. **(C)** Co-immunoprecipitation analysis was used to validate the interaction between gp120 and TNFR2. The IgG group was included as a control to ensure the specificity of the experiment. **(D)** Detection of gp120 and TNFR2 binding using bio-layer interferometry, with five concentrations of gp120 (2μM、1μM、 0.5μM、0.25μM、0.125μM) to determine the affinity. *K*_d_ values were determined from the BLI results, providing a quantitative assessment of the equilibrium binding affinity between gp120 and TNFR2. One-way ANOVA was used to analyze group differences. Significance levels: * p < 0.05, ** p < 0.01, *** p < 0.001, **** p < 0.0001, ns no significant difference.

For further verification, co-immunoprecipitation (Co-IP) assay was performed to study protein-protein interactions between TNFR2 and gp120. To this end, gp120 was incubate with lysates of TNFR2-overexprssing cells, followed by pulling down the protein complexes with gp120-specific antibody which conjugated magnetic beads. The results showed that TNFR2 was present in the immunoprecipitate, indicative of the existence of a direct physical interaction between gp120 and TNFR2 (Fig.1C). The binding of gp120 to TNFR2 was further evaluated using bio-layer interferometry (BLI) to measure biomolecular interactions in real time. Biotinylated human TNFR2 protein was immobilized on a streptavidin (SA) sensor tip and exposed to a gradient of gp120 concentrations, ranging from 125 nM to 2 µM. With 5 µg/ml of immobilized TNFR2-Biotin, gp120 could bind to TNFR2 with high affinity (Fig.1D). As the concentrations of gp120 increased, the response signal enhanced correspondingly, indicating specific and concentration-dependent binding to TNFR2. BLI analysis of the binding kinetics revealed a affinity constant (KD) of 86.5 nM for the gp120- TNFR2 interaction (Fig.1E), indicative of a high affinity and a strong, specific binding interaction.

### TNFR2 interrupts the binding of gp120 to CD4

The entry of HIV-1 into cells was relied on interaction of gp120 with CD4 and CCR5 or CXCR4. In addition to these receptors, several studies have shown that HIV-1 can also bind to other receptors on cell surface(21, 22). Although these receptors exhibit high affinity for gp120, most of them do not affect the entry of HIV-1. These receptors primarily function as attachment factors, retaining viral particles on the cell surface without interfering with the viral infection process. We thus further examined the effect of TNFR2 on the interaction between gp120 and CD4. To visualize the competition between TNFR2 and CD4 for binding to gp120, AlphaFold3 was used to predict the complex structure of TNFR2 and gp120(17). The resulting model exhibited high confidence in the overall complex structure, providing a detailed and reliable representation of the interaction (Fig.2A). The model revealed that TNFR2 bound to gp120 at a specific epitope that is critical for the interaction between CD4 and gp120. The interaction of surface between gp120 and TNFR2 was analyzed in detail, showing that it buries a total area of 1,139.5 Å². The structural analysis reveals a specific interaction interface between TNFR2 and gp120, involving several key amino acid residues. On the TNFR2 side, residues W62, Y65, Y68, Q72, L73, and V75 contribute to the binding, while on gp120, residues L71, T72, T74, M426, W427, and Y435 are involved. These residues form a compact interface likely stabilized by hydrophobic contacts—such as between W62 and W427, and L73 and L71—as well as potential hydrogen bonds, particularly involving Q72 and T72. This substantial interaction surface involves both electrostatic and hydrophobic interactions, with key residues forming hydrogen bonds, salt bridges, and hydrophobic contacts that stabilize the gp120-TNFR2 complex (Fig.2B-2C). Additionally, a comparative analysis of the binding epitopes revealed that the TNFR2 binding epitope on gp120 aligns well with the CD4 binding epitope (Fig.2D). This overlap suggests that the binding of TNFR2 to gp120 could sterically hinder the binding of CD4, and may interfering with the CD4-mediated entry mechanism of HIV-1. Thus, the structural alignment indicates that TNFR2 and CD4 likely compete for the same binding site on gp120.

**Figure 2.**
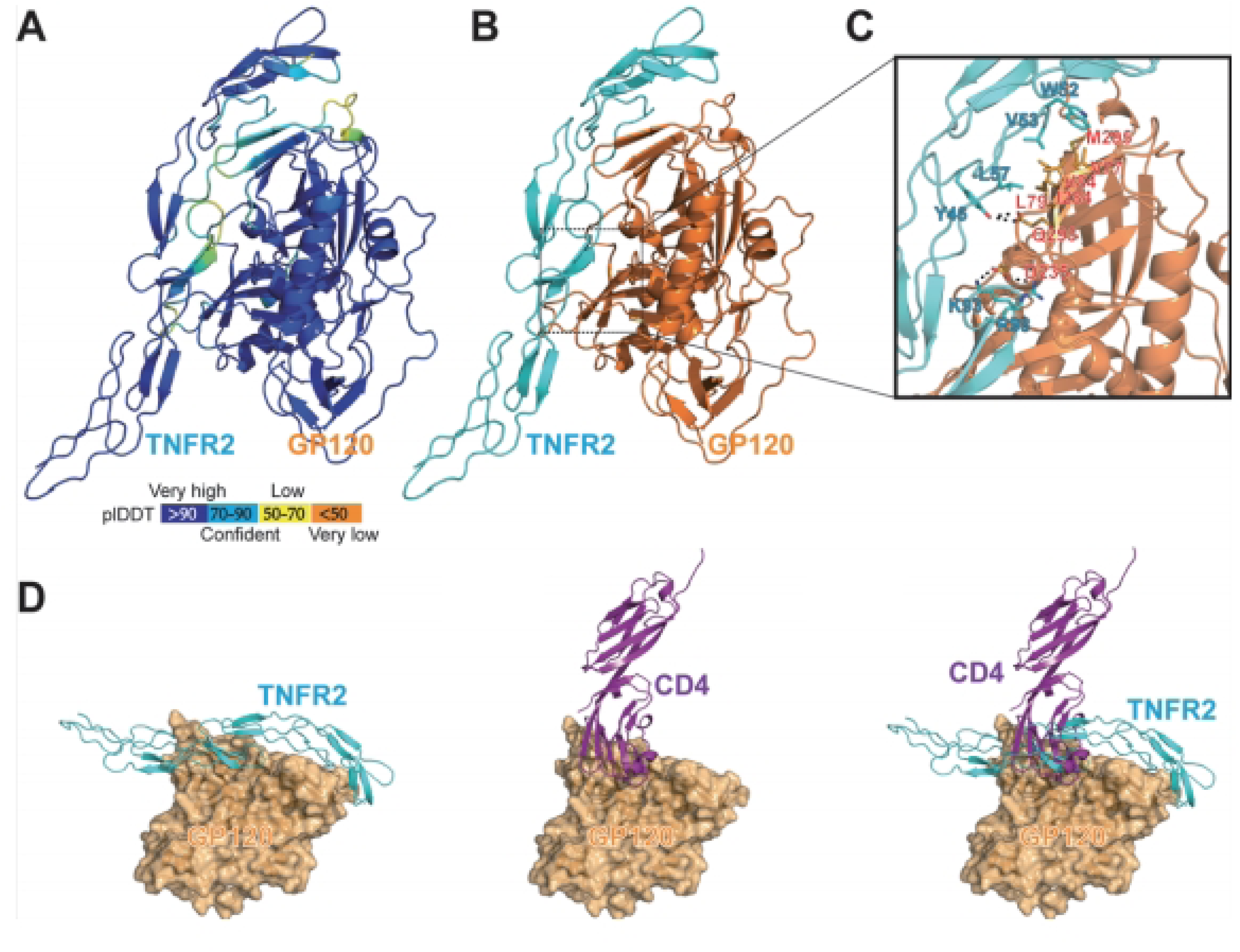
The TNFR2 epitope on gp120 aligns well with the CD4 binding epitope. **(A)** Evaluation of TNFR2–gp120 binding site interactions of Alpha Fold 3 structures. pLDDT as a measure of conformational plasticity. **(B)** The interactions of TNFR2 (blue) and gp120(orange). **(C)** Hydrogen bonds between residues in the TNFR2- gp120 complexes. **(D)** Comparison of the binding epitopes of TNFR2 and CD4 on gp120.

To verify this hypothesis, we investigated the competitive binding between TNFR2 and CD4 for binding to gp120. CD4 was immobilized on the surface of a plate and then incubated with gp120 in the presence or absence of TNFR2 protein. Remarkably, the addition of TNFR2 protein led to a significant reduction in gp120 binding to CD4, with an inhibition rate reaching up to 70% (Fig.3A). This finding demonstrates that TNFR2 competes with CD4 for binding to gp120 which is supportive to our hypothesis. To further verify it at the cellular level, we conducted experiments using Jurkat cell line, a commonly used cell model in the study of HIV-1 infection(23). The binding of gp120 protein to Jurkat cells was quantified by detection of the His tag on the gp120 protein. The result of analysis showed that in the absence of TNFR2, approximately 75% of the cells exhibited gp120 binding. In contrast, the addition of TNFR2 protein resulted in the 65% reduction of gp120 binding to the cells, e.g., only ∼10% of cells exhibited gp120 association (Fig.3B-3D). To validate the specificity of this inhibitory effect, we used an unrelated control protein in parallel. No inhibition of gp120 binding was observed with this control, supporting that the blocking effect is specific to TNFR2 (Supplementary Fig 1B). The inhibition of gp120 binding to the cells by TNFR2 appeared to be dose-dependent, and IC50 was 106.9nM (Fig.3E-3F). And almost 200 nM TNFR2 can basically inhibit the binding of gp120 to Jurkat cells to the greatest extent (Figure 3E).

**Figure 3.**
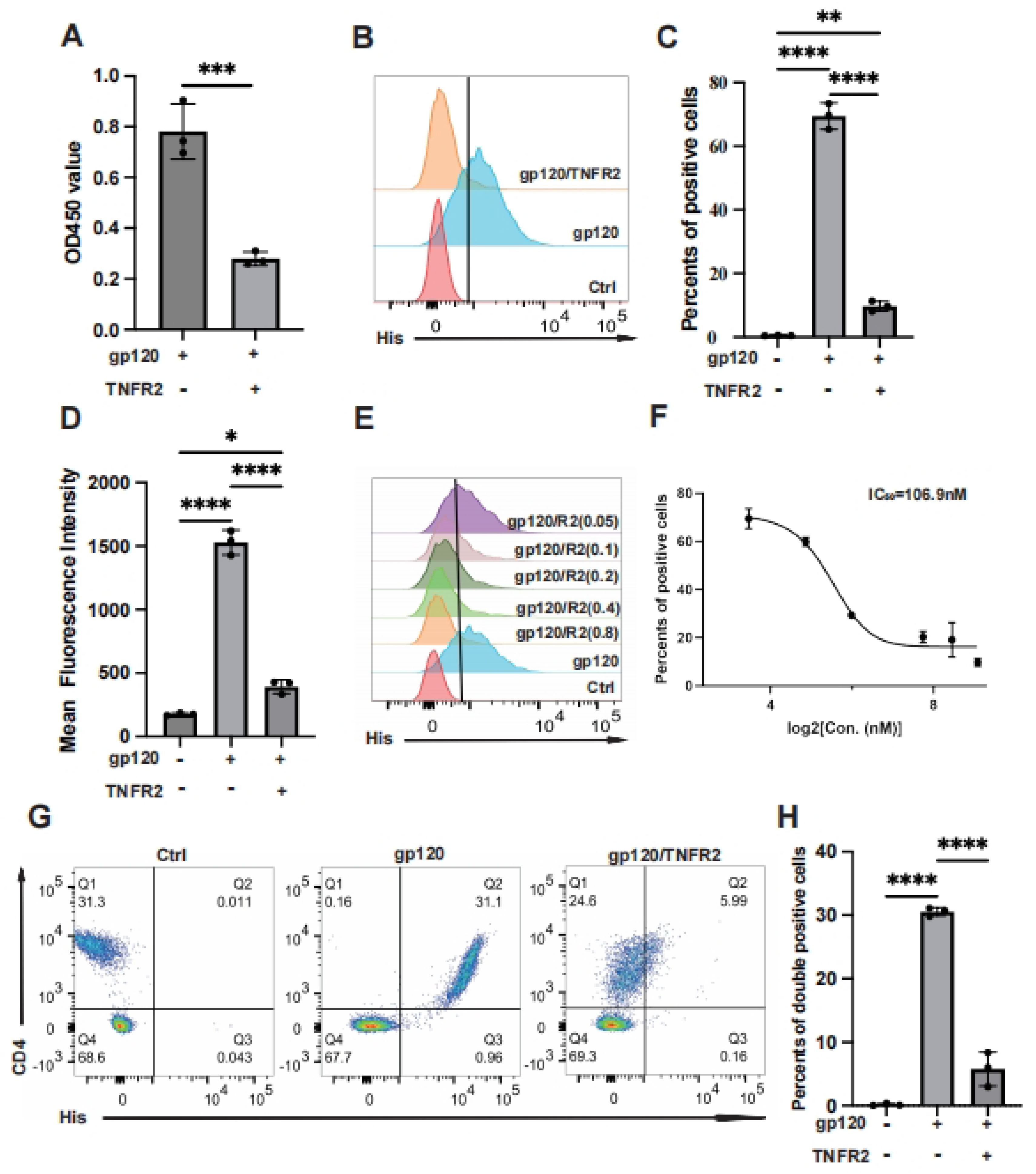
The TNFR2 protein blocks the binding of gp120 to CD4. **(A)** ELISA was used to assess whether TNFR2(2µM) can inhibit the interaction between gp120 and immobilized CD4 on the plate. The final OD450 measurement statistics are shown. **(B)** Flow cytometry was utilized to evaluate cell-bound His-tag, indicating the binding of gp120-His to the Jurkat cells, and also assessed if TNFR2 (1µM) can block gp120-cell binding. **(C)** The percentage of positive cells determined from flow cytometry analysis results. **(D)** The statistical analysis of mean fluorescence intensity. **(E)** Further examination assessed whether the blocking effect of TNFR2 (50nM 、 100nM 、200nM、400nM、800nM) exhibited dose-dependency. **(F)** The half-maximal inhibitory concentration (IC50) was calculated through the dose-dependent inhibition curve. **(G)** Flow cytometry was utilized to detect the binding of gp120 to human peripheral blood mononuclear cells and to evaluate the effect of TNFR2. **(H)** The statistical analysis of the double positive cells. One-way ANOVA was used to analyze group differences. Significance levels: * p < 0.05, ** p < 0.01, *** p < 0.001, **** p < 0.0001, ns no significant difference.

To verify this the in primary human cells, PBMCs from healthy donors were treated with gp120, with or without TNFR2. We could confirm that gp120 exclusively bound to CD4 cells in PBMCs (Fig.3G), as shown with FACS analysis (Supplementary Fig.2A-2B). The binding of gp120 to the cells were potently inhibited by TNFR2, and 200nM of TNFR2 resulted in 80% inhibition of gp120 binding (Fig. 3G-H). Together, our data demonstrate that TNFR2 could potently block the binding of gp120 to CD4 at both the cellular and molecular levels.

### TNFR2 inhibits HIV pseudovirus infection in CD4+CCR5+ cells

Following our earlier observations, we aimed to determine whether TNFR2 could inhibit the cellular infection of HIV-1 by blocking the binding of gp120 to CD4. To explore this possibility, we utilized a pseudovirus-based assay system. This system employed HIV-1 enveloped pseudoviruses that were engineered to express the gp120 envelope protein and encapsulate genes encoding green fluorescent protein (GFP). This design closely mimics the structure and behavior of authentic HIV-1 viruses while lacking the viral genome, making them safe and reliable tools for laboratory investigations.

For this experiment, HEK293T cells co-expressing CD4 and CCR5 were exposed to these pseudoviruses. The expression of green fluorescence was monitored as a measurable indicator of viral infection efficiency, providing a quantitative approach to assess infection rates. Remarkably, we observed that treatment with exogenous TNFR2 protein at a concentration of 1 µM led to a substantial reduction in the infection rate of HIV-1 pseudoviruses. Specifically, the infection rate was reduced by 20% compared to untreated control cells (Fig. 4A-4B). Furthermore, this inhibitory effect of TNFR2 on pseudovirus infection demonstrated a dose-dependent relationship, with increasing concentrations of TNFR2 resulting in progressively greater reductions in infection efficiency (Fig. 4C-4D). These findings indicate that TNFR2 has the capacity to mitigate HIV-1 pseudovirus infection by functionally blocking the binding of gp120 to CD4. This conclusion is further supported by the observed reduction in infection efficiency with higher doses of TNFR2, emphasizing its role as a competitive inhibitor in this context (Fig. 4E). To further explore the antiviral role of TNFR2 in T cells, we repeated the infection experiments using Jurkat cells overexpressing CCR5 as the target cells. Consistent with our findings in HEK293T-CD4-CCR5 cells, pre-treatment of the virus with TNFR2 protein also led to a clear reduction in infection efficiency in these CCR5⁺ Jurkat cells, supporting the broader inhibitory effect of TNFR2 on HIV-1 entry across different cell types (Fig. 4F-4G).

**Figure 4.**
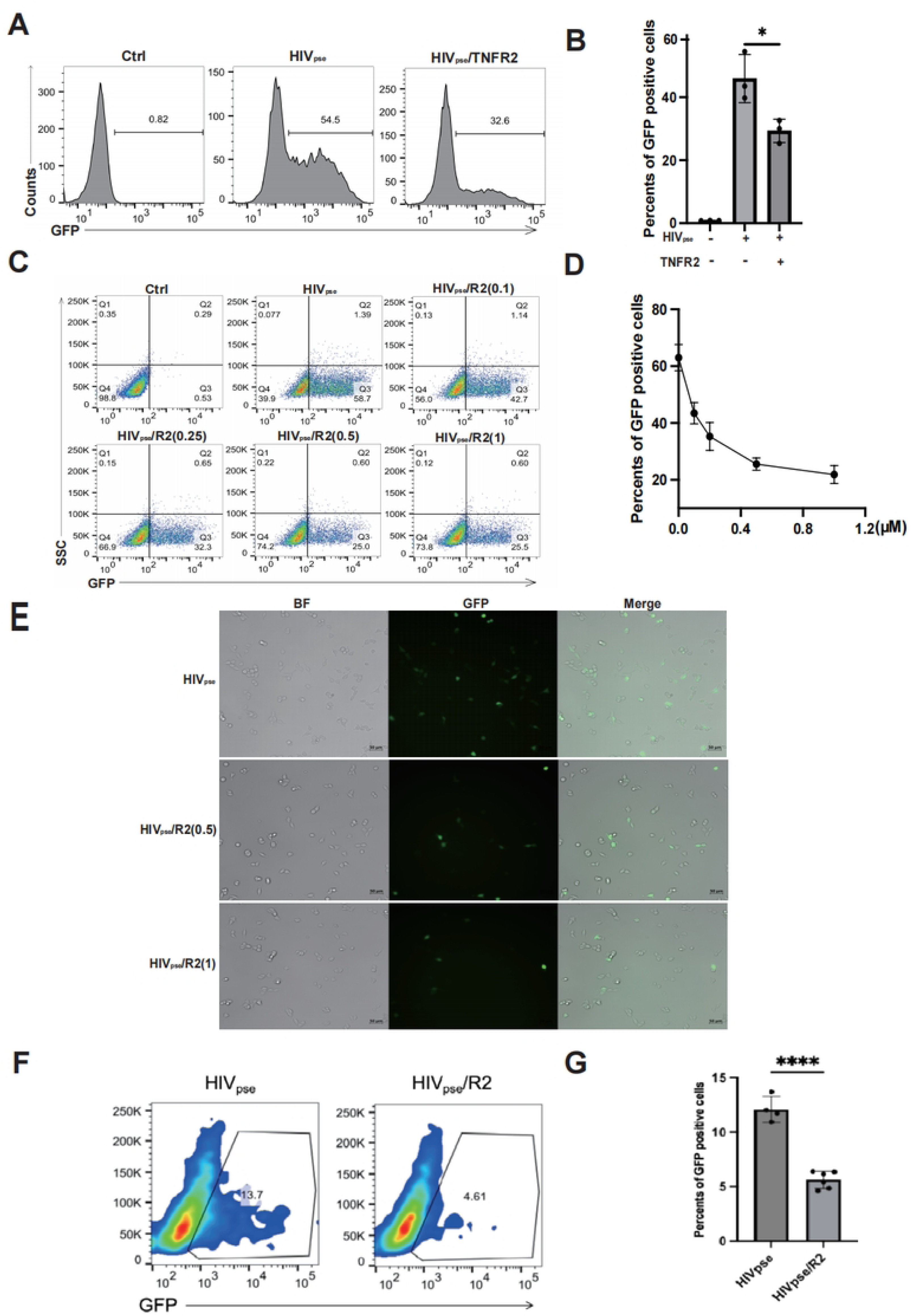
The TNFR2 protein can inhibit HIV pseudovirus infection of CD4+CCR5+ cells. **(A)** HIV pseudovirus carrying green fluorescent protein genes were used to infect CD4+CCR5+293T cells (MOI=10). The flow cytometry was used to analyze the rate of GFP-positive cells to illustrate the virus infection efficiency. **(B)** The statistical analysis for the ratio of GFP positive cells in each group. **(C)** Further examination assessed whether the blocking effect of TNFR2 (0.1µM、0.25µM、0.5µM、1µM) exhibited dose-dependency. **(D)** The statistical analysis for the dose-dependent effects. **(E)** Infected CD4+CCR5+ 293T cells were observed by fluorescence microscopy on the 12h post infection. **(F)** Flow cytometry analysis of Jurkat-CCR5 cells infected with HIV-1 pseudovirus, with or without pre-incubation of the virus with 10 µg/mL TNFR2 protein. **(G)** Quantification of infection efficiency. One-way ANOVA was used to analyze group differences. Significance levels: * p < 0.05, ** p < 0.01, *** p < 0.001, **** p < 0.0001, ns no significant difference.

Collectively, these results suggest a promising potential for TNFR2 in therapeutic strategies targeting the interaction between gp120 and CD4 to prevent HIV-1 infection.

### Membrane-bound TNFR2 expression inhibits HIV-1 pseudovirus infection

After establishing that soluble TNFR2 protein can inhibit HIV-1 infection by blocking the interaction between gp120 and CD4, we next investigated whether membrane-bound TNFR2 expression on host cells could similarly modulate viral susceptibility. We employed a Jurkat cell model overexpressing CCR5 and partially expressing TNFR2. Following HIV-1 pseudovirus infection, flow cytometry analysis revealed that TNFR2⁺ Jurkat cells exhibited a 1.5-fold reduction in GFP-positive cells compared to the TNFR2⁻ subsets (Fig.5A-5B). These results provide direct evidence that membrane-bound TNFR2 negatively regulates HIV-1 infection, reinforcing its role as a protective factor in cells.

**Figure 5.**
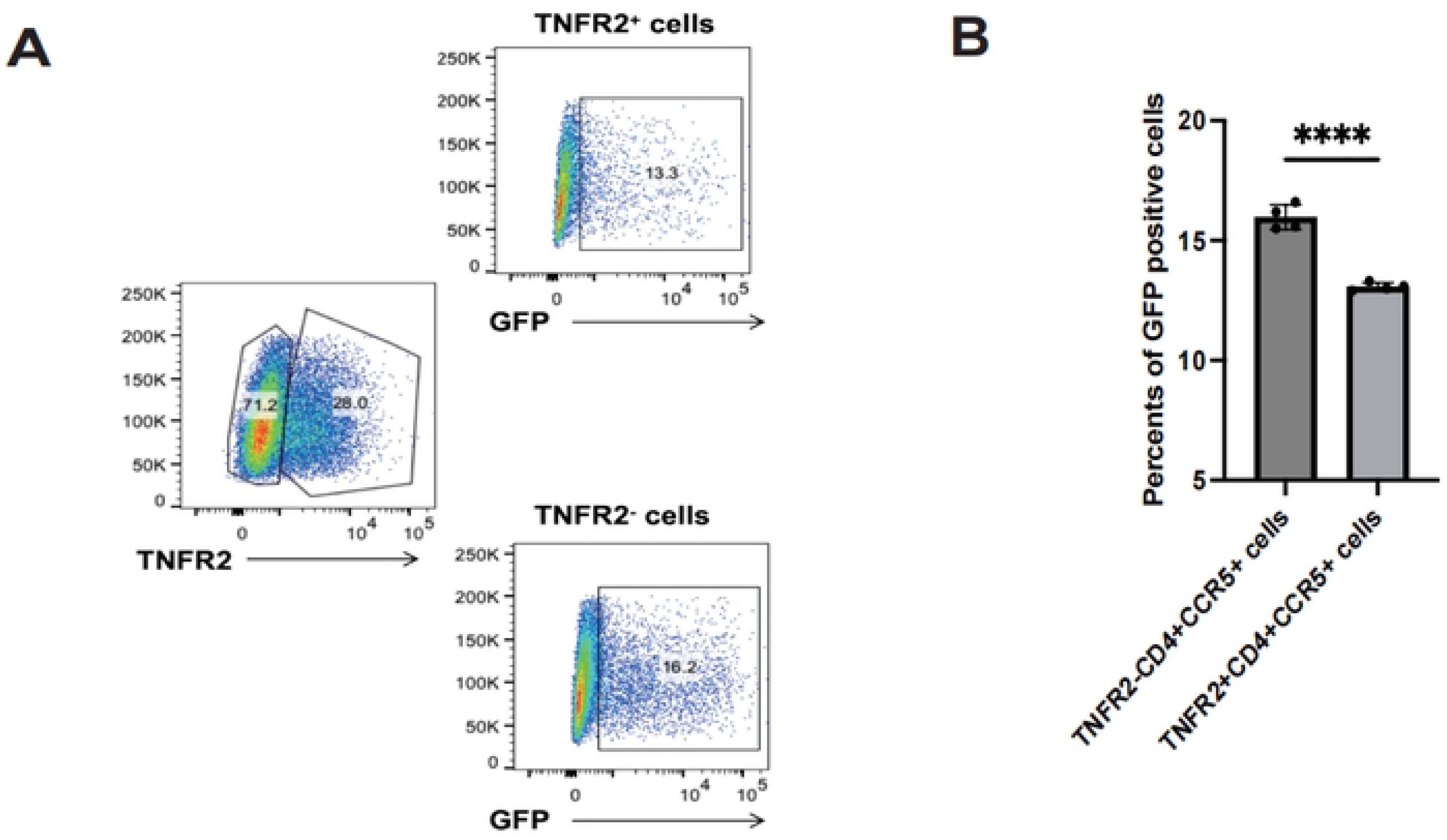
TNFR2 expression inversely correlates with HIV-1 pseudovirus infection. **(A)** CCR5⁺ Jurkat cells with partial TNFR2 overexpression (∼30% TNFR2⁺) were infected with GFP-expressing HIV-1 pseudovirus. GFP expression was analyzed in TNFR2⁺ and TNFR2⁻ subpopulations by flow cytometry. **(B)** Quantification of the percentage of GFP⁺ cells in TNFR2⁺ and TNFR2⁻ populations to evaluate pseudovirus infection efficiency. Significance levels: * p < 0.05, ** p < 0.01, *** p < 0.001, **** p < 0.0001, ns no significant difference.

### TNFR1 affects the binding of gp120 to CD4 but promotes HIV-1 infection

Due to the structural similarities between TNFR1 and TNFR2, as well as their shared ligand TNF, we extended our investigation to evaluate whether TNFR1 interacts with gp120. We first performed ELISA assays and confirmed that recombinant TNFR1 protein directly binds to gp120 in a concentration-dependent manner, similar to TNFR2 (Fig.6A). Specifically, when Jurkat cells were treated with TNFR1, we observed a significant reduction in the binding of gp120 to these cells (Fig.6B-6C). A similar trend was observed in primary lymphocytes, where the presence of TNFR1 markedly reduced gp120 binding (Fig.6D-6E).

**Figure 6.**
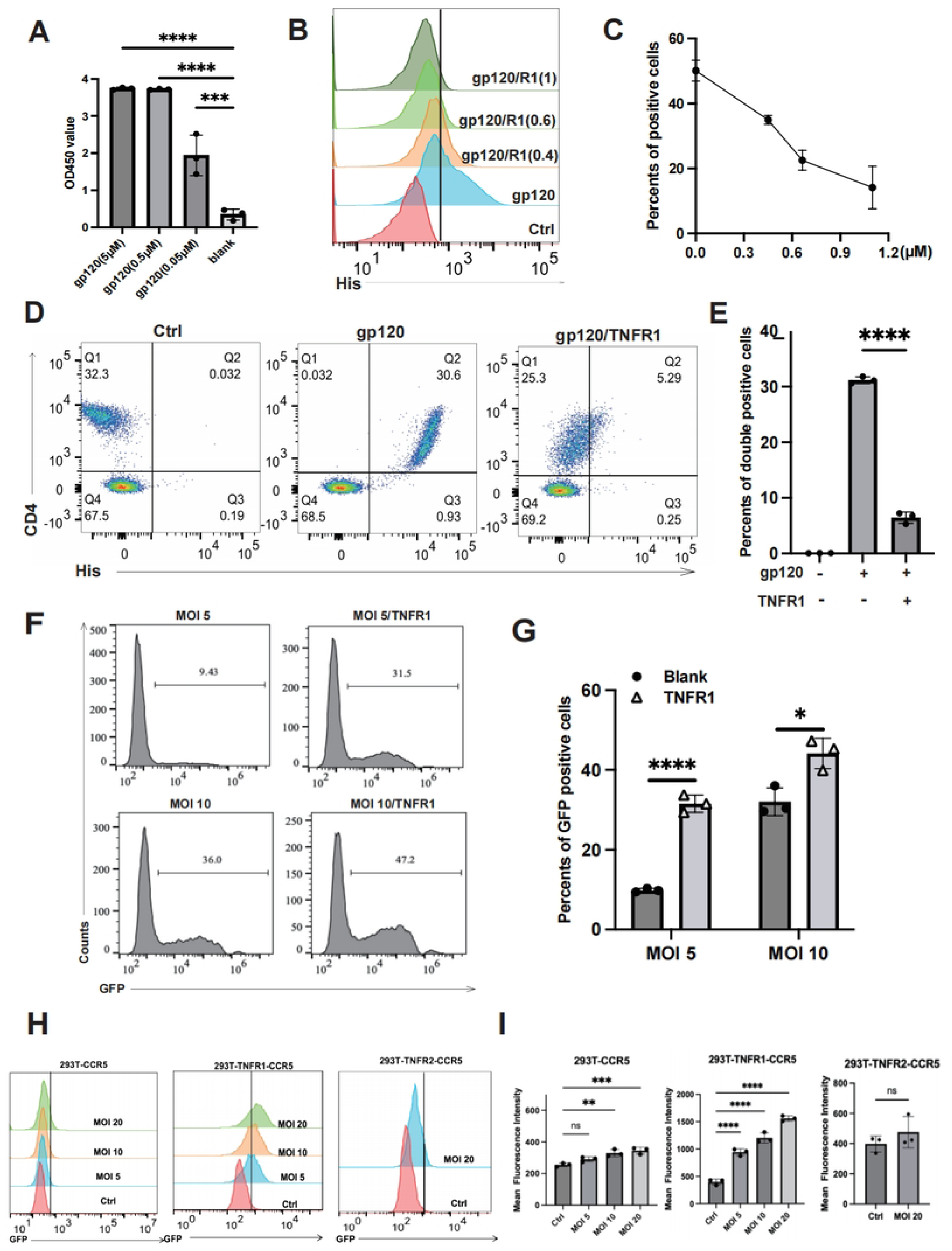
TNFR1 impacts the binding of gp120 to CD4 but promotes the virus infection. **(A)** The binding interaction between gp120 and TNFR1 was assessed using an ELISA assay with varying concentrations of gp120 (5μM、0.5μM、0.05μM、 blank). **(B)** Flow cytometry was utilized to evaluate cell-bound His-tag, indicating the binding of gp120 to the Jurkat cells, and also assess if TNFR1 (0.4µM、0.6µM、 1µM) can block gp120-cell binding. **(C)** The percentage of positive cells determined from flow cytometry analysis results. **(D)** Flow cytometry was utilized to detect the binding of gp120 to human peripheral blood mononuclear cells and to evaluate the blocking effect of TNFR1. **(E)** The statistical analysis of the double positive cells. **(F)** HIV pseudovirus carrying green fluorescent protein genes were used to infect CD4+CCR5+293T cells (MOI=5、MOI=10). The flow cytometry was used to analyze the rate of GFP-positive cells to illustrate the virus infection efficiency. **(G)** The percentage of GFP positive cells determined from flow cytometry analysis results. **(H)** CD4⁻CCR5⁺ 293T, TNFR1^+^CD4⁻CCR5⁺ 293T, and TNFR2^+^CD4⁻CCR5⁺ 293T cells were infected with GFP-expressing HIV-1 pseudovirus. Infection efficiency was assessed by flow cytometry based on the percentage of GFP⁺ cells. **(I)** Quantification of infection efficiency. Significance levels: * p < 0.05, ** p < 0.01, *** p < 0.001, **** p < 0.0001, ns no significant difference.

Interestingly, when we further investigated the role of TNFR1 in the context of HIV-1 infection, we uncovered a surprising contrast. While we initially hypothesized that TNFR1, like TNFR2, might act to inhibit HIV-1 infection by blocking the gp120-CD4 interaction, our results showed the opposite. In experiments using HIV-1 enveloped pseudoviruses, TNFR1 was found to facilitate viral infection rather than inhibit it (Fig.6F-6G). This unexpected finding suggests that TNFR1 may play a distinct role in promoting the entry or replication of HIV-1 pseudoviruses. These results highlight a fundamental difference in the functional roles of TNFR1 and TNFR2, with TNFR1 potentially enhancing HIV-1 infectivity under certain conditions.

To explore the underlying mechanism of this enhancement, we further examined whether TNFR1 could facilitate viral infection in the absence of CD4. We employed engineered cell lines expressing CCR5 alone (as a control), or co-expressing CCR5 with either TNFR1 or TNFR2 (Fig.6H-6I). Notably, TNFR1—but not TNFR2—was able to promote HIV-1 pseudovirus infection under these CD4-deficient conditions. These results suggest that TNFR1 may act as a cofactor by inducing conformational changes in gp120 that expose or stabilize CCR5-binding sites, thereby enhancing viral entry even without canonical CD4 engagement.

## DISCUSSION

In this study, we present compelling evidence that the TNFR2 protein plays a crucial role in modulating the entry of HIV-1 into CD4 cells, serving as an effective inhibitor of viral infection. Our findings suggest that this inhibitory effect is primarily mediated through the competitive binding between TNFR2 and CD4 to the HIV-1 envelope protein gp120, which is essential for viral entry. By binding to gp120, TNFR2 outcompetes the CD4, preventing the necessary conformational changes in gp120 required for its subsequent interaction with the coreceptor CCR5. This disruption in the viral entry process effectively blocks the fusion of the virus with the host cell membrane, thereby impeding HIV-1 infection and reducing viral replication within the target cells. The mechanistic underpinning of this process is complex but intriguing. It appears that TNFR2 acts not merely as a passive competitor for gp120 binding but as a dynamic regulator of viral entry. By preventing the interaction between gp120 and CD4, TNFR2 interferes with the early stages of HIV-1 infection. This disruption in the viral lifecycle is a critical step in preventing HIV-1 from successfully establishing infection in immune cells.

Our findings are consistent with prior studies demonstrating the impact of TNF mutant proteins on HIV-1 infection. Specifically, pretreating macrophage cells with a TNF mutant protein that selectively binds to TNFR2 resulted in a significant reduction in HIV-1 virus entry into the cells(11). We believe that their findings are based on TNF mutants activating TNFR2 signaling, leading to upregulation of TNFR2 expression on the cell surface, thereby inhibiting HIV-1 infection. We propose that the therapeutic potential of these TNF mutants lies not only in their ability to bind TNFR2 but in their capacity to activate TNFR2 signaling, leading to an upregulation of TNFR2 expression on the cell surface. The increased availability of TNFR2 on the cell membrane could, in turn, create an environment that is less permissive to HIV-1 infection, as the heightened receptor density disrupts the viral entry process.

Our previous studies have demonstrated that TNFR2 is highly expressed on Tregs and plays an important role in maintaining their immunosuppressive function(14). Tregs, known for their essential role in immune homeostasis, rely on TNFR2 signaling for their survival and suppressive activity(24–26). When the HIV virus approaches Treg cells, TNFR2 acts as a binding partner for the virus, potentially preventing viral entry and mitigating the impact of infection on these critical immune regulators. This interaction suggests a mechanism that limits HIV’s ability to compromise Tregs, thereby preserving immune suppressive function.

In this study, we found that both TNFR1 and TNFR2 are capable of competitively binding to the HIV-1 envelope protein gp120, thereby interfering with its interaction with the CD4 receptor. This competitive binding may block the initial step of viral attachment to host cells and provides a novel mechanistic insight into the regulation of HIV-1 entry. Previous studies have shown that the structure of gp120 is highly dependent on its binding state, and certain proteins or molecules can induce conformational changes in gp120 that affect viral fusion and entry(27). Interestingly, TNFR1 and TNFR2 exhibited markedly different effects on HIV-1 pseudovirus infection. TNFR2 showed a strong inhibitory effect, while TNFR1 appeared to promote infection. These contrasting outcomes may be attributed to the distinct conformational changes induced in gp120 upon binding with TNFR1 or TNFR2. It has also been reported that gp120 can expose or conceal different functional domains—such as the V3 loop or fusion peptide—depending on the nature of its interacting partner, thus affecting the virus’s infectivity (28). Based on our observations, we speculate that TNFR2 binds to a region of gp120 that sterically hinders CD4 attachment or induces an unfavorable conformation for subsequent fusion events, thereby blocking viral entry. In contrast, TNFR1 may bind to a different epitope or interact with gp120 in a way that partially exposes fusion-relevant structures, inadvertently facilitating viral entry under certain conditions. These findings not only highlight the unique roles of TNFR1 and TNFR2 in the context of HIV infection but also provide a theoretical basis for developing antiviral strategies that leverage the inhibitory potential of TNFR2.

In conclusion, our study provides crucial insights into the role of TNF receptors in the dynamics of HIV-1 infection, shedding light on the complex interplay between HIV viral entry and host immune responses. Although biosafety constraints prevented us from conducting live virus experiments, our pseudovirus-based assays still provided strong theoretical support for these findings. We demonstrate that TNFR2 plays a critical and unique role in inhibiting HIV-1 entry into host cells, offering a potential mechanism by which the immune system can limit HIV viral propagation.

Specifically, our research strongly supports the design of anti-HIV-1 drugs by developing TNFR2 derivatives. By modifying the TNFR2 protein, we aim to enhance the bodies resistance to HIV-1, thereby effectively curbing viral invasion and replication. This approach not only reduces viral proliferation within host cells but also increases patients’ immunity, promising a new way for future HIV-1 infection treatments. In summary, our study provides a fresh perspective on understanding the mechanisms of HIV-1 infection. By elucidating the functions of TNF receptors, we hope to make meaningful contributions in tackling the global challenge of HIV/AIDS.

## DATA AVAILABILITY STATEMENT

The data underlying figures are available in the published article and its online supplemental material.

## ACKNOWLEDGMENTS

**Funding** This project was funded by Macau Science and Technology Development Fund (0007/2022/AKP and 0099/2021/A2), University of Macau (MYRG2022-00260-ICMS, CPG2024-00031-ICMS, MYRG-GRG2024-00300-ICMS), State Key Laboratory of Quality Research in Chinese Medicine (University of Macau)(SKL-QRCM (UM)-2023-2025), Applied Research Programs of Guangdong-Hong Kong-Macao Innovation Center sponsored by Guangzhou Development District (EF032/ICMS-CX/2021/RITH), Hong Kong and Macao scientific and technological achievements transformation project in Guangdong (2023A0505030012).

## Authorship

Contribution: XC and HS provided project oversight for experimental design, data management, data analysis and interpretation, and methods development; YG and Zh C helped with workflow development, experiments, data processing, management, and analysis; and all authors wrote the manuscript and provided feedback.

Conflict-of-interest disclosure: The authors declared no potential conflicts of interest with respect to the research, author-ship, and/or publication of this article.

Correspondence: Institute of Chinese Medical Sciences, State Key Laboratory of Quality Research in Chinese Medicine, University of Macau, Macau, 999078, P.R. China.

